# Simultaneous cell traction and growth measurements using light

**DOI:** 10.1101/206649

**Authors:** Shamira Sridharan, Yanfen Li, Louis Foucard, Hassaan Majeed, Basanta Bhaduri, Alex Levine, Kristopher Kilian, Gabriel Popescu

## Abstract

Understanding cell mechanotransduction is important for discerning matrix structure-cell function relationships underlying health and disease. Despite the crucial role of mechanochemical signaling in phenomena such as cell migration, proliferation, and differentiation, measuring the cell-generated forces at the interface with the extracellular matrix during these biological processes remains challenging. An ideal method would provide continuous, non-destructive images of the force field applied by cells, over broad spatial and temporal scales, while simultaneously revealing the cell biological process under investigation. Toward this goal, we present the integration of a new real-time traction stress imaging modality, Hilbert phase dynamometry (HPD), with the technique of spatial light interference microscopy (SLIM) for label free monitoring of cell growth. HPD relies on extracting the displacement field in a deformable substrate, which is chemically patterned with a fluorescent grid. The displacements introduced by the cell are captured by the *phase* of the periodic signal associated with the grid, borrowing concepts from holography. The displacement field is uniquely converted into forces by solving an elasticity inverse problem. Because the measurement of displacement only uses the epi-fluorescence channel of an inverted microscope, we can simultaneously achieve measurements in transmission. We performed SLIM and extracted cell mass on the same field of view in addition to the measured displacement field. We used this technique to study mesenchymal stem cells and found that cells undergoing osteogenesis and adipogenesis exerted larger and more dynamic stresses than their precursor. Our results indicate that the MSCs develop the smallest forces and growth rates. We anticipate that simultaneous cell growth and traction measurements will improve our understanding of mechanotransduction, particularly during dynamic processes where the matrix properties provide context to guide cells towards a physiological or pathological outcome, e.g., tissue morphogenesis, or cancer metastasis.

## INTRODUCTION

Interactions between cells and their microenvironment involve a complex combination of pushing and pulling forces that in turn affect processes such as cell adhesion, migration, proliferation and differentiation ^1^. The cytoskeletal network, comprised of filamentous actin, intermediate filaments, and microtubules, mediate the transmission of forces within the cell ^2^. At the cell membrane, the cytoskeletal elements bind to focal adhesion proteins that mediate interactions with the extra-cellular matrix (ECM) through cell surface integrins. The mechanical forces generated at these interfaces influence biochemical signaling, in a process termed *mechanotransduction*, which has important implications on cell fate, and downstream tissue form and function ^3^. A focal adhesion, thus, serves as both the point at which the cell may exert forces on its surroundings and as the cell’s window into mechanical changes occurring in the surrounding tissue, leading to adaptive changes within the cell. Understanding mechanotransduction is especially critical in cancer cell biology, where processes such as cell lamellipodia extension, migration, and angiogenesis, are mediated through interactions with the surrounding ECM and can dramatically influence the aggressiveness and metastatic potential of the cancer ^4–7^. In addition to pathological processes, during normal development, mechanochemical signals regulate stem cell lineage determination, where the ECM provides context to guide the integration of multiple cues in the microenvironment. Stem cells show a degree of lineage plasticity and can shift their state through interrogation of ECM mechanics and composition. The mechanical interactions between stem cells and their environment influences proliferation, migration and differentiation, towards the regulation of wound healing, tissue morphogenesis and homeostasis ^8–11^.

In recent years, significant progress has been reported in the development of quantitative techniques to measure cell generated forces (see Ref. ^12^ for a recent review). These methods can be broadly classified into two categories: *active,* i.e., based on measuring the response of the cell to application of external forces, and *passive*, built on measuring the substrate deformation due to intrinsic, cell-generated forces. Common methods within the first category are atomic force microscopy, optical tweezers, and magnetic tweezers ^13–16^. However, these methods suffer from the limitations in spatial sampling and restrictive thresholds for the measurable forces. *Traction force microscopy* techniques that have gained the most widespread adoption employ micropatterned pillars, textured substrates, and coated fluorescent beads. Traction forces from micropillar arrays are calculated from the bending of soft pillars of known mechanical properties ^17,18^. However, by restricting the cell adhesion cites to the locations of the micropatterned pillars, this technique is not an ideal representation of 2D cell culture. Substrate deformation measured from textured PDMS substrates are an alternative method ^19^. However, the minimum stiffness with which PDMS can be fabricated is Young’s modulus, ~12kPa, which is stiffer than many in vivo environments. The technique also requires florescent tagging of the focal adhesion sites within the cell. A similar method looks at the displacement of topographically patterned dots, however the method has multiple shortcomings from computational assumptions including limited spatial resolution, and the requirement that one observe single cells, or, at least, cells with high degree of spatial separation ^20^. Another established approach involves the incorporation of fluorescently tagged beads within an elastic substrate to study the traction of multiple cells ^21,22^. However, this method requires the removal of cells from the substrate surface in order to obtain the initial configuration of the incorporated beads, which is a tedious and error-prone process, and also limits the force measurement to a single time point. Recently, *confocal* traction force microscopy has been developed based on nanodrip-printed monocrystalline array of fluorescent marks ^23^. This approach alleviates some of the previous limitations, but at the same time relies on specialized, high-precision substrate preparation and retains the need for tracking fluorescent particles individually.

Here, we present multimodal approach that monitors cell growth using *spatial light interference microscopy* (SLIM) simultaneously with cell generated traction force through a novel technique we refer to as *Hilbert phase dynamometry* (HPD). HPD measures forces exerted by cells in *real-time*, over extended periods of time, without disturbing the culture. The cells are grown on flat, deformable substrates and the in-plane displacement field is continuously measured with high spatial and temporal resolution. Engineering the deformable substrate at subcellular scales with a customized 2D grid, which is fabricated through patterning fluorescent adhesion proteins, enables the adherent cell-induced strain field to be contained in the 2D phase map of the complex analytic signal associated with the periodic grid. The principle of HPD is rooted in the phase reconstruction used in *off-axis holography*, as developed by Leith and Upatnieks in the 1960s’ ^24^ (for a review on phase reconstruction and imaging, see also ^25^). From the displacement field, we solve the *inverse* elasticity problem and extract a traction force vector field. Due to the uniform sampling of the substrate via the fluorescent grid, HPD eliminates the need for tracking individual particles and can be applied continuously, without removing the cells from culture. A valuable feature offered by HPD stems from its compatibility with a conventional microscope. In particular, as the force field is measured in the epi-fluorescence channel, the transmission illumination can be used for simultaneous, complementary measurements. We used spatial light interference microscopy (SLIM)^26–29^ for performing quantitative phase imaging (QPI) on the same field of view as the fluorescence channel. QPI is an emerging field of label-free imaging that has found important applications in biomedicine ^25^. Among them, studying cell growth has perhaps the broadest potential ramifications as it addresses this “long-standing question in biology” ^30^. Thus, we combined the measurements of HPD and SLIM to obtain simultaneous cell growth and traction information during stem cell differentiation. Using mesenchymal stem cells (MSCs) as a model adherent adult stem cell that shows high responsivity to ECM properties^21,31–33^, we found that cells undergoing differentiation, osteogenesis and adipogenesis, exerted larger and more dynamic stresses than their precursor. By using integrated HPD-SLIM we uncover a relationship between MSC generated traction, growth, and differentiation. Our results indicate that the MSCs exert the smallest forces and have the lowest growth rates compared to their differentiated progeny.

## RESULTS

Figure 1 illustrates the principle of HPD. First, the PA gel is chemically activated (see details on the gel preparation in Materials and Methods and Supplemental Information Section S3) and stamped with fluorescent protein, to create a 9 μm period grid in both x and y directions. Briefly, 10kPA polyacrylamide hydrogels were pipetted onto a hydrophobically treated glass slide. Using a pattern master 2D grid with a 9 micron period, adhesion proteins were stamped on the hydrogel (Fig. 1B). The substrate was then uniformly exposed to fibronectin to ensure homogeneous cell adhesion to the substrate (Fig. 1C). In this way FITC conjugated and non-FITC conjugated adhesion proteins are placed in alternating intervals, such that the cells do not sense the grid and, upon traction, generate bending of the substrate in both directions. The cells were seeded and allowed to settle on the substrate (Fig. 1D). The deformations in the substrate were measured from the phase of the 2D periodic signal associated with the grid. Finally, the force field is extracted via a *Cerruti-type inverse problem* (see Fig. 1E and Materials and Methods for details)^34^

**Figure 1.**
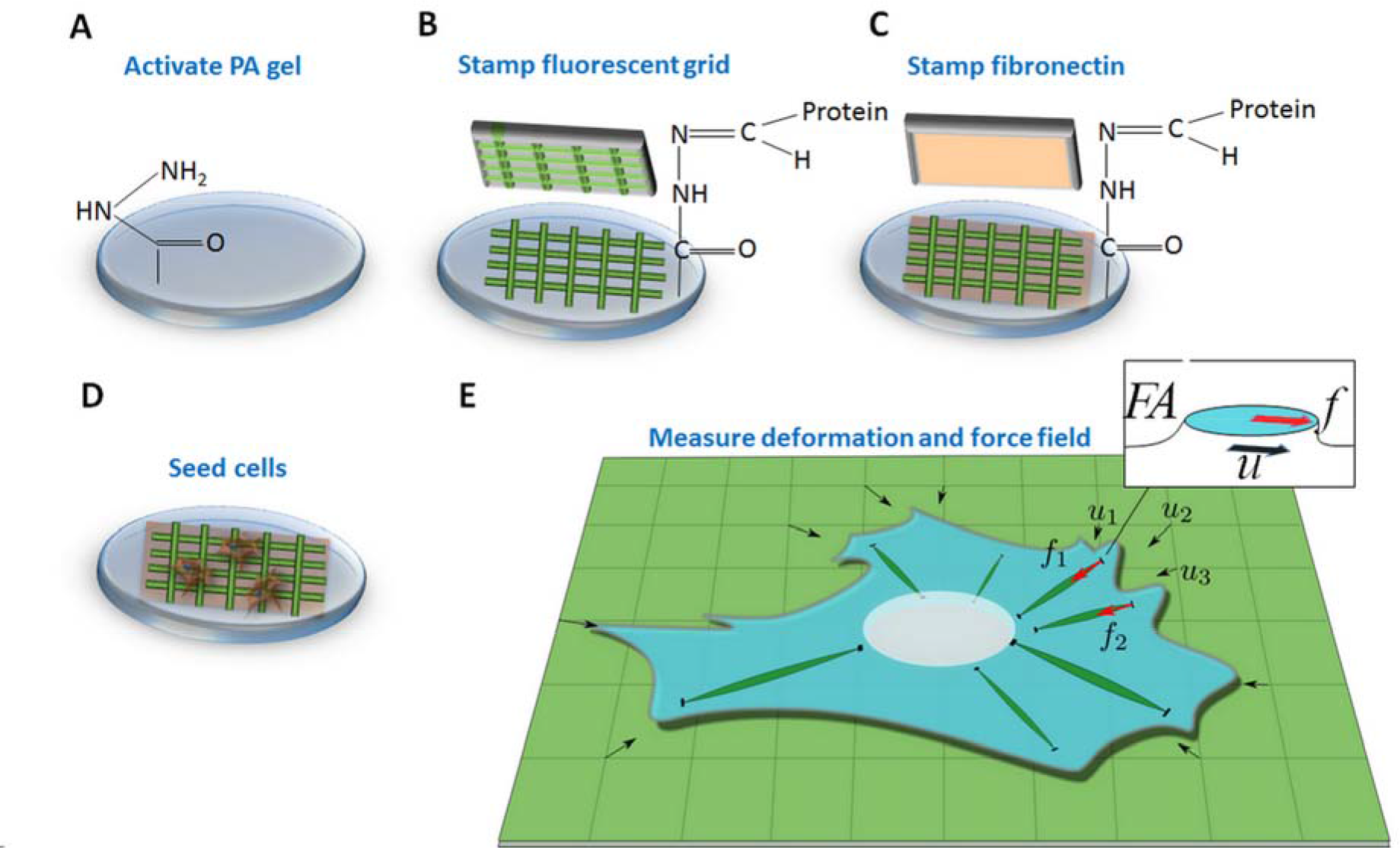
Patterned polyacrylamide hydrogels enables measurements of traction force. (A) Polyacrylamide hydrogel are activated to contain hydrazide groups. (B) A fluorescent grid containing adhesion proteins is stamped onto the activated hydrogel. (C) Remaining regions are filled with non-fluorescent adhesion proteins. (D) Cells are seeded onto the gel and allowed to attach. (E) The traction force f exerted by the cell on the substrate can be retrieved by measuring the resulting substrate deformation u.

The experimental setup and procedure for extracting the displacement field are shown in Figure 2. The measurements were performed with an inverted microscope, outfitted with both a SLIM module (Cell Vista SLIM Pro, Phi Optics, Inc.), and an epi-fluorescence optical train (see Fig. 2A and Materials and Methods for further information on the optics). The epi-fluorescence channel provides images of the grid, while SLIM renders quantitative phase images in transillumination, which can be further analyzed in terms of cell dry mass density ^29^. Note that the live cells are unlabeled and, thus, can be imaged without damage over many hours. Because of their common optical path, the SLIM and fluorescence channels are overlaid at the pixel level without the need for numerical registration.

**Figure 2.**
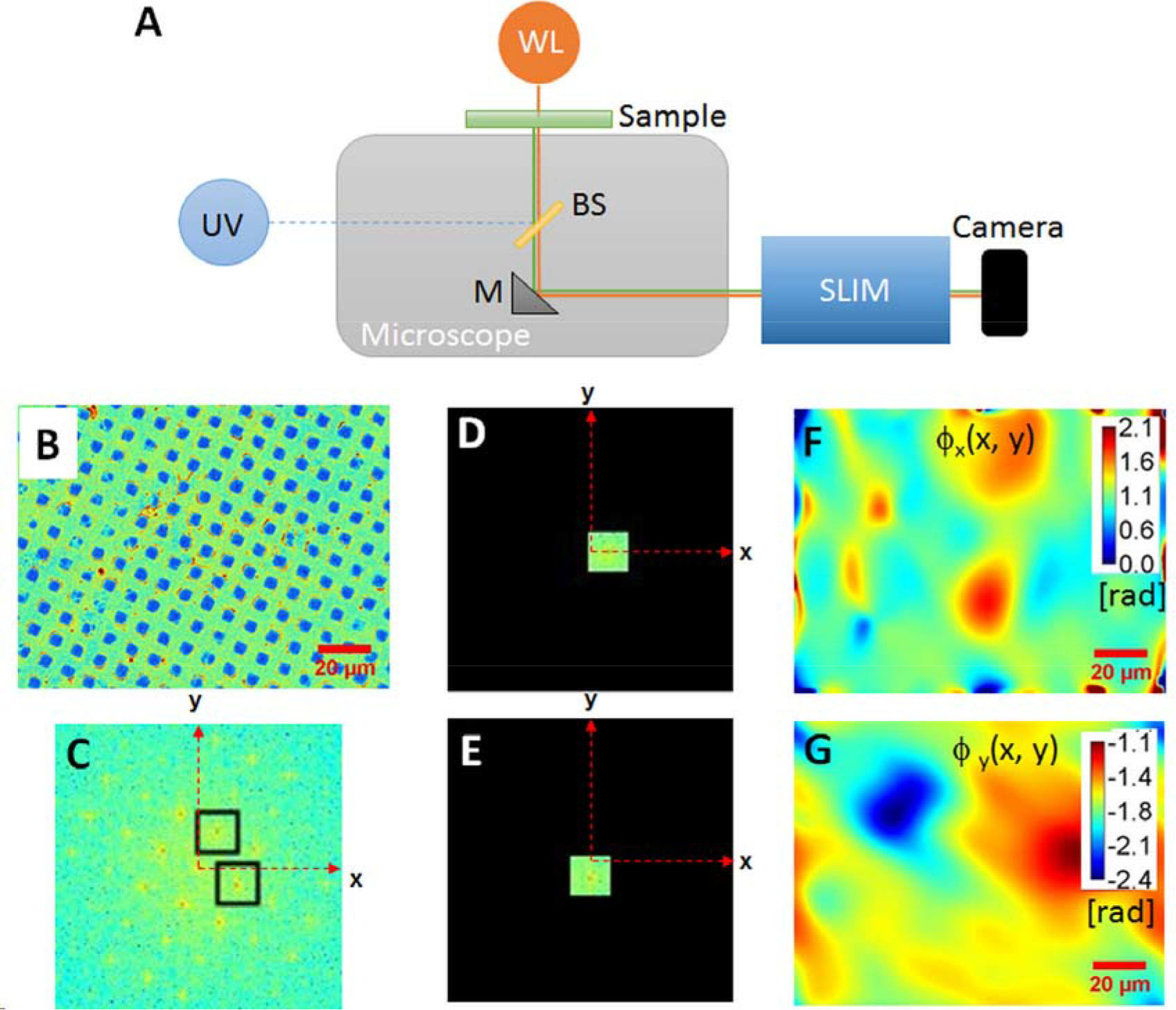
Schematic illustration of the X and Y displacement map reconstruction process. (A) The raw fluorescence image of the grid (B) Fourier transform of the raw image (C) Zooming into the central region of the Fourier transform shows well separated orders due to the periodicity of the grid. (D-E) The spectrum is band pass filtered over the regions shown. (F-G) Taking the inverse Fourier transform of each image in F and G generates the phase shift maps along the x and y axes.

The image of a fluorescence grid is shown in Fig. 2B and the absolute value of its Fourier transform in Fig. 2C. Because the grid is not perfectly sinusoidal in shape, the Fourier transform of its image generates multiple orders along each direction. However, if we only retain the first orders in both *x* and *y* directions, the analysis is equivalent to that of a perfect sinusoidal grid. The signal of interest along each direction has the form (See Supplemental Section 1)

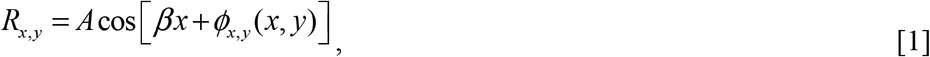

where R_x,y_ are the sinusoidal fluorescence intensities (real-valued signals) along x and y, *ϕ*_x,y_(*x*,*y*) the respective phases that incorporate the displacement information, and *β* the spatial frequency of the grid, *β*=2*π/9 rad/μm*. We apply a spatial frequency filter that selects the first order in the *x* and *y* direction as shown, respectively, in Figs. 2D and 2E. Inverse Fourier transforming the signals in Figs. 2D and 2E results in complex signals, namely the *complex analytic signals* associated with R_x,y_. The concept of the complex analytic signal associated with a real optical field was exploited early on by Gabor^35^ and served as foundation for his development of holography^36^. These two complex signals, one for each direction, are derived from the grid image via the following expressions

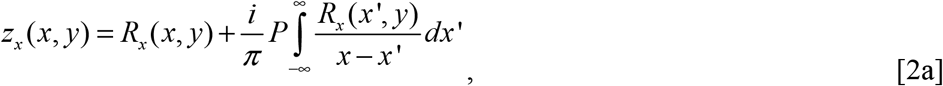

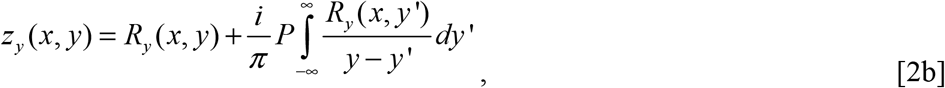

where P stands for principal value integral. The sequence of Fourier transforming the real signal, spatially filtering, and Fourier transforming back to the spatial domain amounts to a Hilbert transform ^37^, which is captured in the HPD acronym. The argument of each signal provides the deformation of the grid at each point in the field of view along each direction, *ϕ*_*x*,*y*_ = arg(*z*_*x*,*y*_).

The phase maps associated with the grid in Fig. 2B are shown in Figs. 2F and 2G, respectively. This phase information, in radians, is converted into spatial displacement, *u*, in microns, by noting that 2π radians corresponds to a displacement of a grid period, namely,

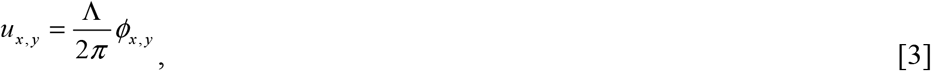

where Λ is the period, Λ = 9 *μm*.

The microscope can switch between the fluorescence and SLIM channels in 0.7 s, which makes it particularly appealing for studying cell growth and traction simultaneously. The instrument is completely programmable and can scan large fields of view in x-y, acquire z-stacks, as well as time lapses over scales from fractions of seconds to many days. We cultured bone marrow derived mesenchymal stem cells (MSCs), and then subjected them to media containing soluble supplements supporting adipogenesis or osteogenesis for 1 week (see Materials and Methods for details on the differentiation process). We chose to explore MSCs because these cells are a promising cell type for autologous therapy, and no technique to date has been able to relate cell-matrix traction and growth during lineage specification. Figure 3 demonstrates that displacement and dry mass density maps can be obtained simultaneously and quantitatively by our method.

**Figure 3.**
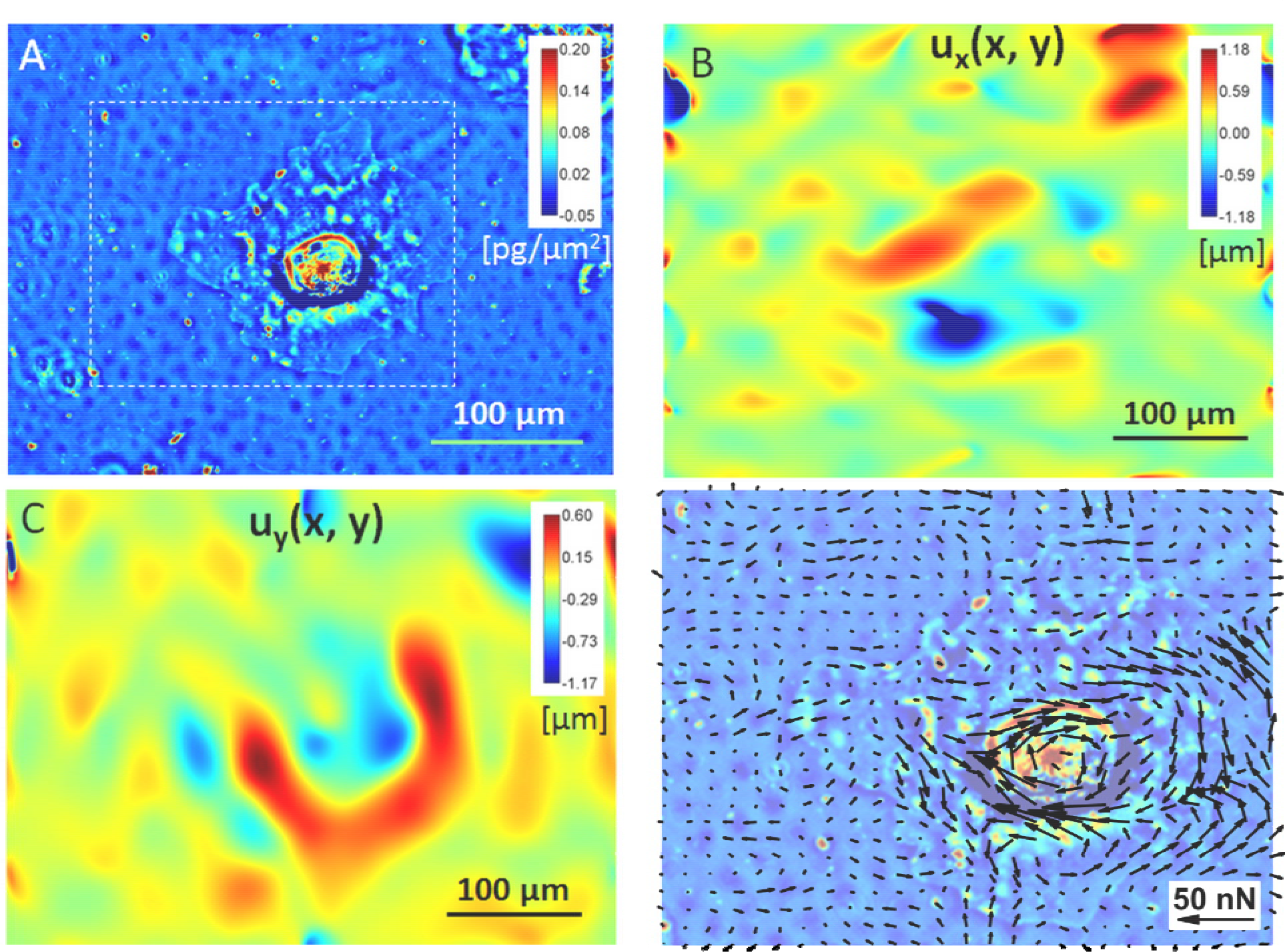
Force field calculations from displacement maps. (A) Quantitative phase image of adipocytes on polyacrylamide hydrogel coated with 2D fluorescent fibronectin grid at 9μm periodicity at the beginning of the experiment. (B) X displacement map calculated from the phase map of the fluorescent grid deformations. (C) Y displacement map calculated from the phase map of the fluorescent grid deformations. (D) Cell traction force calculated from the X, Y displacement maps are overlaid on the phase image of the adipocyte.

In order to extract the force field from the measured displacement maps, we solve a linear elasticity inverse problem (see Supplemental Information Section 2 for details). Essentially, due to the linearity of the problem, the 2D displacement field **u** at position **x** and a disk distributed force of density **f** applied at an arbitrary position ***x’*** are related via a simple matrix vector multiplication:

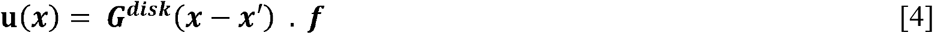

In Eq. 4, the 2×2 matrix ***G***^*disk*^ is a response (or transfer) function that describes the displacement response of the substrate, and is obtained by solving a Cerruti type problem ^34^ (see Supplemental Section 2 for details on the derivation). We chose to use disk distributed forces here to represent the traction applied by the cell at focal adhesion sites. If one were given the locations of all focal adhesions and the tractions that they produce, the surface displacements are then found by computing the sum of the displacements associated with each focal adhesion separately, computed using Eq. (4). Given a measured displacement map of size N × N, and a grid of hypothetical FAs position of size M × M (with M ≤ N), the problem of finding the forces exerted by the cell on these FAs can be written as the following linear system:

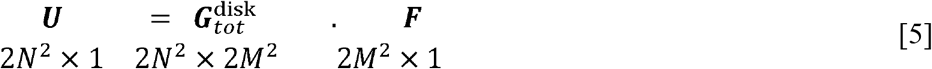

In Eq. 5, 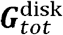 represents the summed response functions between multiple displacements and forces position. Finally, the force field ***F*** is computed by inverting the system above using a least square approximation, which concludes the HPD procedure. Figure 3D shows an overlay of dry mass and force field obtained by HPD. Supplemental movies S1-6 show the real-time measurement of the displacement force fields for several cells.

Previous reports have shown changes in cytoskeleton structure and contractility in MSCs undergoing osteogenesis or adipogenesis ^38^, which could lead to changes in focal adhesions and traction force ^39^. In order to test the sensitivity of our system, we seeded patterned hydrogels with MSCs that were exposed to either basal, osteogenic, or adipogenic media for one to two weeks. To confirm differentiation of MSCs, cells were stained after one week in differentiation media with Oil Red O to confirm adipogenesis and with alkaline phosphatase to confirm osteogenesis. Figure 4 illustrates the concomitant measurements of traction and SLIM acquired over 12 hours, with a temporal sampling of 15 minutes. There are clear morphological differences between the three cell types. In particular, the MSCs stand out due to their significantly smaller size. The overlays between SLIM and the magnitude of the force field clearly show that the forces applied by the MSCs are the smallest. The results also indicate that the forces exerted by the cells become stronger with time.

**Figure 4.**
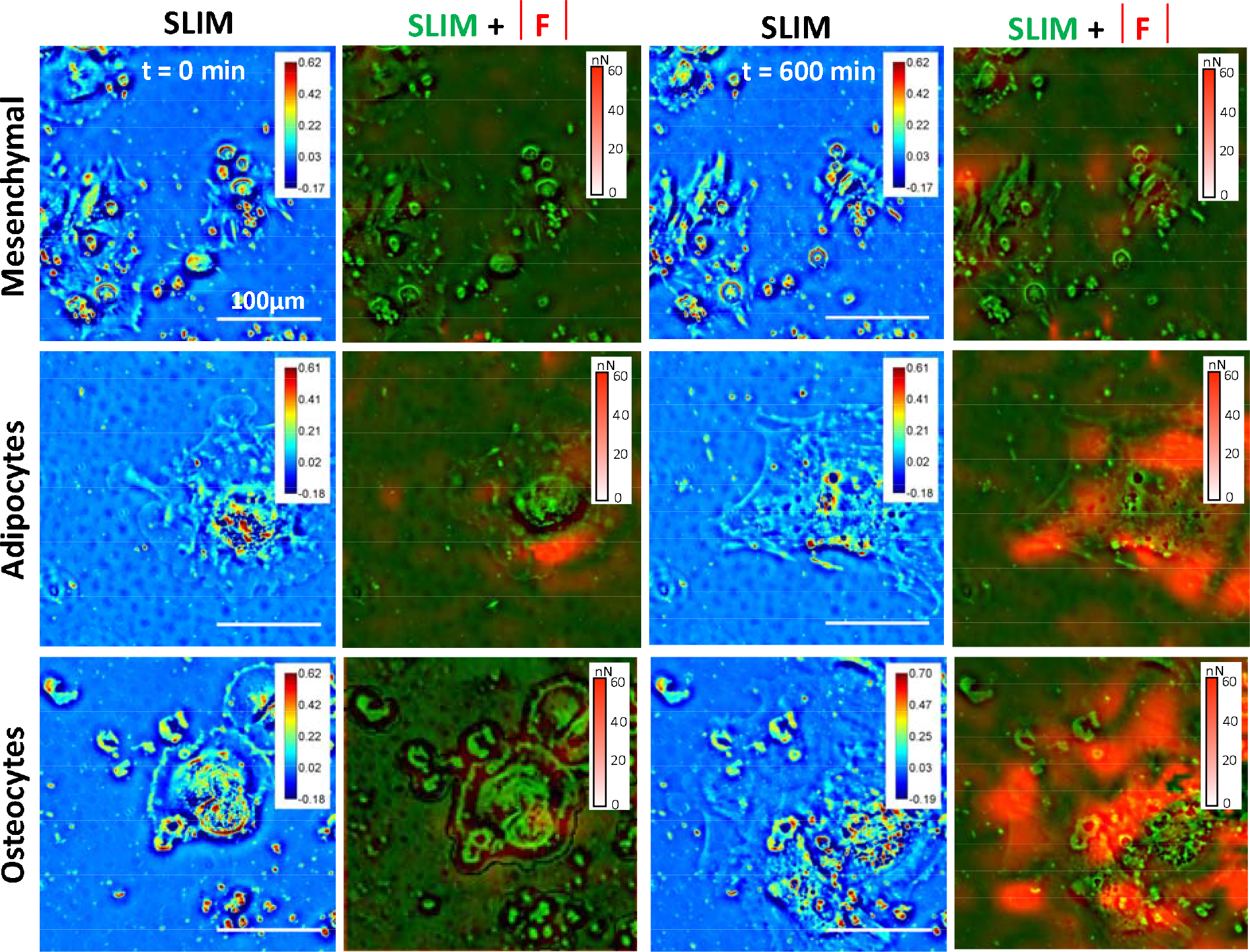
Cell growth and force measurements. Screenshots of mesenchymal (row A), adipocytes (row B) and osteocytes cells (row C) taken at time t=0 (left column), t = 100 (middle column) and t = 600 min (right column). Each time step shows the SLIM image, and the combined SLIM (green scale map) and force magnitude (red scale map).

Histological staining of MSCs exposed to the different media formulations demonstrates the appearance of alkaline phosphatase in the osteogenic conditions and accumulation of lipid droplets in the adipogenic conditions (Figure 5A). The histogram of all the measured forces (Figure 5B) indicate that the mesenchymal stem cells apply the lowest mean force and also display the narrowest spread in force magnitude (22.9±17.1nN). The largest mean tractions are produced by the adipocytes (51.5±39.2nN) followed by cells undergoing osteogenesis. At the same time, the lowest dry mass growth was shown by the MSCs and the highest by the osteoblasts. Previous work has demonstrated increased traction stress exerted by cells on microposts during the initial stages of adipogenesis and osteogenesis compared to MSCs ^39^, thus supporting our observations. Furthermore, using our combined approach of SLIM and HPD, we are able to simultaneously measure changes in cell mass and dynamic traction in real time during the initial stages of lineage specification. This advance provides the first technique where dynamic interactions of cells and their matrix can be queried during cell and tissue level processes in situ. Supplemental movies S7-S9 illustrate the real-time overlay.

**Figure 5.**
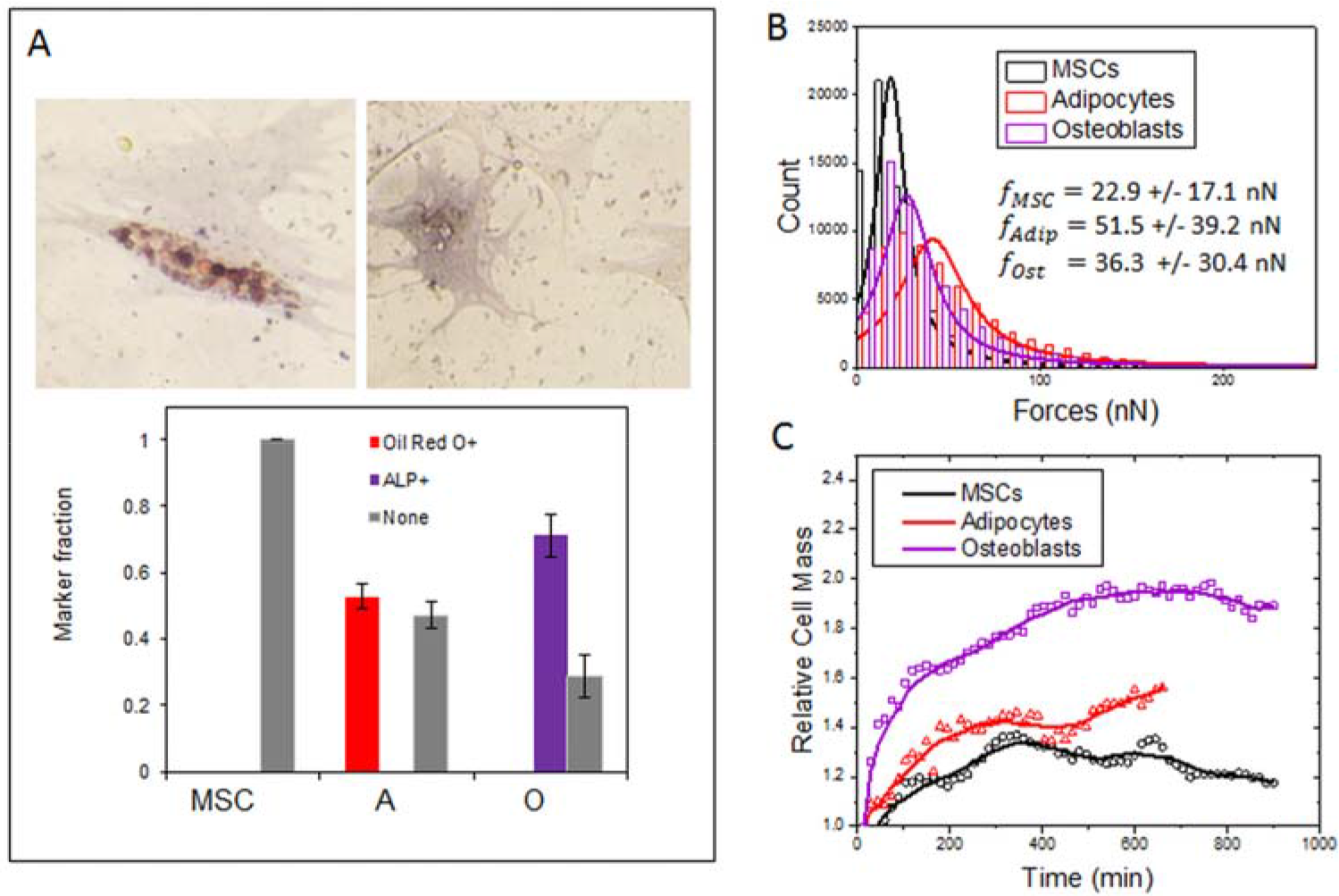
Cell lineage specific traction force and growth calculations. (A) At the end of the experiment, the cells were stained with Oil Red O+, an adipogenic marker, and ALP+, a marker for osteogenesis. The marker fractions were used to verify lineage-specific cell differentiation.(B) Histograms showing the distributions of cell traction forces associated with mesenchymal stem cells, adipocytes, and osteoblasts, across the entire experiment. The average magnitude of force exerted by adipocytes was the highest and that of mesenchymal stem cells was the lowest.Growth curves displaying the relative cell mass of mesenchymal stem cells, adipocytes and osteoblasts across time. Osteoblasts have the highest growth rate.

### Summary and discussion

Using beaded polyacrylamide hydrogels for traction force microscopy, it has been recently demonstrated that MSCs exert the highest traction stress on hydrogels conjugated with fibronectin compared to laminin and collagen ^21,40^. Therefore, we conjugated fibronectin in patterned grids to explore cellular traction using HPD and cell growth with SLIM techniques, for MSCs after exposure to media supplements that promote differentiation to osteoblast or adipocyte lineages. Cells exposed to osteogenic supplements, including ascorbic acid, β-glycerophosphate, and dexamethasone, exerted higher traction stress over time compared to MSCs cultured under standard growth media. Cells exposed to adipogenic supplements, including indomethacin, insulin, dexamethasone, and isobutylmethylxanthine, exerted significantly higher traction stress compared to both MSCs in growth media and those undergoing osteogenesis. These results show some consistency with work by Fu et al. that demonstrated enhanced traction force from cells undergoing differentiation ^39^. However, while they observed an initial spike followed by a rapid decay to basal levels in traction force for cells undergoing adipogenesis (over 7 days; micropost arrays), here we observe higher average traction from cells undergoing adipogenesis compared to osteogenesis and basal conditions. These differences may be related to experimental conditions, where in our work we trypsinized MSCs after 1 week in differentiating media, followed by transfer to our patterned substrates where the HPD and SLIM techniques were performed under normal media conditions. We believe that performing our analysis in the absence of hormones may aid the unambiguous assessment of cell generated force as a function of cell state. We speculate that our observed differences in cell traction force is related to the evolution of mechanosensing machinery and the actomyosin network that has previously been shown to accompany specification to adipocyte and osteoblast lineages ^41^.

Previous studies of MSC proliferation during differentiation has demonstrated enhanced division for cells undergoing differentiation, with the supplements dexamethasone and ascorbic acid playing clear roles ^42,43^. Here we demonstrate the power of our integrated multimodal imaging approach by simultaneously quantitating traction force and a measure of cell growth by SLIM. Consistent with known relationships between differentiation and proliferation, we observe enhanced cell dry mass growth with adipogenesis > osteogenesis > basal conditions. Multipotent MSCs are known to show a degree of quiescence with low division rates when cultured in niche-mimetic conditions ^32,43,44^ Using the combined tools of HPD and SLIM enable the in situ tracking of relationships between extracellular recognition, force transduction, and specific bioactivities including growth and differentiation.

These techniques can be readily coupled with live cell fluorescent reporters, or fluorescent in situ hybridization techniques, in the same setup to further enable high-content data acquisition for deciphering how cell generated force corresponds to biological outcomes in real time. Furthermore, the generality of the hydrogel fabrication approach will allow virtually any combination of stiffness, and protein, proteoglycan or appropriately tagged synthetic biorecognition motif, to be gridded across the substrate to explore mechanotransduction in a model cell biological system of interest.

## Materials & Methods

### Gel Preparation

10kPA polyacryamide hydrogels were fabricated by mixing 5% polyacrylamide (Sigma Aldrich) and 0.15% bis-acylamide (Sigma Aldrich) as previously described (Tse & Engler, 2010). 0.1% Ammonium Persulfate (APS, Sigma Aldrich) and 0.1% Tetramethylenediamine (TEMED, Sigma Aldrich) was added to the acrylamide mixture and pipetted onto a hydrophobically treated glass slide (Fisher). An amino-silanized glass cover slip was then flipped onto the solution and allowed to incubate for 20 minutes. Gels were then lifted off and immersed in 55% hydrazine hydrate (Fisher) for two hours to convert amide groups to reactive hydrazide groups followed by 5% glacial acetic acid for one hour. Gel stiffness was confirmed with AFM as previously described ^45^. In order to prepare the substrates for protein patterning and cell adhesion, hydrazine hydrate was used to convert PAAm amide groups to reactive hydrazide groups allowing for the conjugation of ECM proteins via coupling of formed aldehyde groups after oxidation with sodium periodate (Fig. 1, Step 1).

### Gel Patterning

A patterned master of photoresist (SU-8, Microchem) was created via UV light through a laser printed mask. Polydimethysiloxane (PDMS, Polysciences, Inc) was then polymerized on top of the master to create a stamp with 9 μm spaced grids, such that FITC conjugated and non-FITC conjugated adhesion proteins were placed in alternating 9 μm intervals along both the X and Y axes. Note that there is no limitation in principle for how fine a grid can be stamped, provided it is resolved by the imaging system. A mixture of 25μg/ml of fibronectin and 25 μg/ml of FITC conjugated fibrinogen were incubated with Sodium Periodate (Sigma Aldrich) for 20 min to yield free aldehydes. This incubation took place on top of the patterned PDMS stamp for 30 mins, dried under air and then applied to the surface of the hydrogel, which had been dried in room temperature for 40 minutes (Fig. 1, Step 2). Next, 25 μg/ml fibronectin on a blank PDMS stamp was applied onto the hydrogel, following the same procedures as the previous step (Fig. 1, Step 3). By following these procedures, we obtained a uniform distribution of adhesion proteins on the gel surface with periodic regions displaying fluorescence signal. The coverslip with the gel was then glued to the bottom of a glass-bottom cell culture dish (MaTek) at two points using tissue adhesion glue (Liquid bandage, CVS).

### Cell Culture and Staining

Human mesenchymal stem cells (MSC, Lonza) were allowed to grow until they reached 70% confluency and then seeded onto a 6-well plate to initiate differentiation processes. The cells were cultured in MSC growth media (low glucose DMEM, 10% FBS, 5% Pen/Strep, Gibco), adipogenic media (Lonza), or osteogenic media (Lonza) for one week. Adipogenic media was rotated between induction and maintenance every 3 days. Cells were then lifted off the substrate with 0.25% trypsin (Sigma Aldrich) and seeded onto glass-bottom dishes (Fig. 1, Step 4) and imaged using the multi-modal SLIM system. At the end of imaging, the cells were fixed with 4% paraformaldehyde (PFA) for 20 minutes and incubated in 60% isopropanol for 5 min followed by immersion in Oil Red O working solution (3:2; 300 mg/mL Oil Red O in isopropanol:DI water, Sigma Aldrich) for 10 min and then BCIP/NBT (Sigma Aldrich) for 10min.

### Multi-modal SLIM/ Fluorescence Imaging System

Spatial light interference microscopy (Cell Vista SLIM Pro, Phi Optics, Inc.) is a QPI system that operates as an add-on module to an existing commercial phase contrast microscope ^26,46^. The back focal plane of the phase contrast objective is projected onto a liquid crystal phase modulator, where programmable phase rings introduce 3 additional phase shifts, in increments of π/2, between the scattered and un-scattered light transmitted through the sample. The phase is computed in real-time using the corresponding intensity images. Using software developed inhouse, the imaging modality can be switched between phase and various fluorescence channels. Thus, we are able to obtain quantitative phase and FITC images of the same field of view.

Mesenchymal stem cells, adipocytes and osteocytes placed on deformable substrates with fluorescent protein grids were imaged using the phase and FITC module on the SLIM system using a 20X/0.45NA objective. The cells were imaged for 12 hours at 15 minute intervals. Typically, 6-8 fields of view were selected from each plate for imaging. The FITC image was taken at the plane of focus for the protein stamp, while the SLIM data were recorded as z-stacks with 2 frames above and below the plane of focus. This insures that the dry mass of the cells is integrated longitudinally, along the entire thickness of the cell. The maximum was projected through the SLIM z-stack, to account for changes in cell structure through the imaging time. The cell mass was calculated from the phase image using the following relationship, as described in detail in ^47,48^:

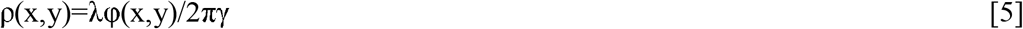

where λ is the central wavelength of the light source, φ(x,y) is the phase value of the corresponding pixel and γ = 0.2ml/g, is the refractive index increment of protein ^49^. Since the mass from multiple cells are averaged for analysis, the mass of each cell at each time point was calculated relative to the first time point. With this procedure, we eliminate the chance that a few heavy cells dominate the mean and growth trends.

## AKNOWLEGEMENT

This work was supported by National Science Foundation (NSF) Grants CBET-0939511 STC, DBI 14-50962 EAGER, IIP-1353368 (to G.P.), DMR-1309188 (to A.J.L.), 1454616 CAR (to K.K.), and National Institutes of Health, HL12175 (to K.K.). Y.L. was supported by the National Science Foundation Graduate Research Fellowship Program under Grant No. DGE – 1144245.

